# Relationship between tumor grade and geometrical complexity in prostate cancer

**DOI:** 10.1101/015016

**Authors:** J.A. Llanos-Pérez, E. Tejera-Puente, E. Izquierdo-Kulich, J.A. Betancourt Mar, M. Nistal, P. González-Peramato, M. Royuela-García, J.M. Nieto-Villar, M. P. De Miguel

**Author notes:** Corresponding author. Author address for correspondence and reprint: Maria P De Miguel, Cell Engineering Laboratory, IdiPaz, La Paz Hospital Research Institute, Paseo Castellana 261, Madrid 28046, Spain.

## Abstract

Prostate cancer exhibits high mathematical complexity due to the disruption of tissue architecture. An important part of the diagnostic of prostate tumor samples is the histological evaluation of cellular and glandular organization. The Gleason grade and score, a commonly used prognostic indicator of patient outcome, is based on the match of glandular architectural patterns with standard patterns. Unfortunately, the subjective nature of visual grading leads to variations in scoring by different pathologists. We proposed the fractal dimension of the lumen and the Lempel-Zip complexity of the histopathological patterns as useful descriptors aiding pathologist to standardize histological classification and thus prognosis and therapy planning.

**Highlights:**

- geometrical complexity of prostate cancer

## 1. Introduction

Cancer is a generic name given to a group of malignant cells which have lost their tissue-specific specialization and control over normal growth. These groups of malignant cells are nonlinear dynamic systems which self-organize in time and space, far from thermodynamic equilibrium [1], and exhibit high complexity [2,3], robustness [4,5] and adaptability [6].

As reported by the WHO in its bulletin No. 297, 2013, cancer is the leading cause of death worldwide. According to the American Cancer Society in 2013 of those diagnosed from with prostate cancer the 12% die due to cancer, and 1 in 6 men are diagnosed with this disease. [7].

When signs of prostate cancer appear, although there are many specific features [8], the clinical aggressiveness of this tumor type is diagnosed through Gleason grade and score, which is based on the observation of different sections of the biopsy or the surgical specimen. Each histopathological picture is assigned a Gleason grade (from 0 to 5 increasingly different from normal), which depends on the architecture of tumor glands and the relationship of the gland with the surrounding stroma (Fig. 1). As the prostate gland and prostate tumors are morphologically heterogeneous, the Gleason score is obtained by the sum of the most predominant Gleason grade and the second most predominant one in frequency [10,11]. In this regard, it has been observed that the architecture of the prostate gland becomes irregular and complicated with increasing grade of agressiveness of the same. [10, 11].

**Figure 1.**
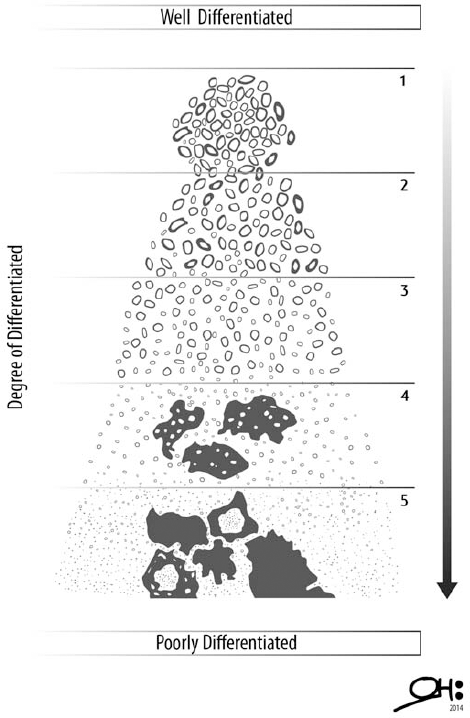
Gleason classification in which the gland structure corresponds to different Gleason grades of prostate cancer. Adapted from [11].

Due to the complexity of the histopathological images, the use of the theory of complex systems, in particular fractal geometry has been proved useful to discern objectively between these complex patterns [12, 13, 14, 15]. The complexity analysis enables to understand the phenomena that determines the evolution of the tumor and can be used as a tool to develop strategies for therapies [12, 16, 17]. In previous studies we have used the fractal dimension to characterize patterns in cervical cancer [18] and found a mathematical relationship between the rate of proliferation of the tumor and its fractal dimension.

Although over the last decade various methods for the analysis of histopathological images of prostate have been developed [15, 17, 19, 20, 21], yet the issue of the complexity of these images and their quantification remains a topical issue and a necessity for the clinical analysis of the same.

The aim of this work is the application of complex systems theory to quantify the agressiveness of prostate cancer, in particular the determination of the capacity fractal dimension (*D_f_*) and the complexity of Lempel-Ziv ( *LZ*) of histopathological images this tumor, which can be used as a complement in the diagnosis and prognosis.

## 2. Materials and methods

### 2.1 Prostate tissue specimen procurement

Data were obtained from the archives of the Pathology Services Prince of Asturias Hospital, Alcala de Henares, Madrid, Spain. All pathological, clinical or personal data were anonymized and separated from any personal identifiers. This study was made with the consent of the patients’ relatives or their family in autopsy cases and from surgical biopsies used for diagnosis. All the procedures were examined and approved by the University of Alcalá and Principe de Asturias Hospital Ethics Committees (reference number SAF2007-61928) and were in accordance with the ethical standards of the Committee for Human Experimentation, with the Helsinki Declaration of 1975 (as revised in Tokyo 2004) and the Committee on Publication Ethics (COPE) guidelines.

Samples used were sections 5 µm-thick, stained with hematoxylin-eosin and observed under an optical microscope (Olympus, Vanox-T, Mod AH-2). In total, 277 images were taken (30 Healthy Tissue, 5 Gleason-1 (G1), 12 Gleason-2 (G2), 76 Gleason-3 (G3), 15 Gleason-4 (G4) and 6 images of Gleason-5). Images were classified by pathologists M. Nistal and P. González Peramato, following guidelines of the consensus of the International Society of Urological Pathology of 2005 [10] and tha Gleason grade 4 modification of 2010 [11]. All images were taken at 40X magnification and a resolution of 1296×972 pixels.

### 2.2 Morphological characterization and Mathematical processing

The morphological characterization of each image was held by software ImageJ 1.43 [22]. All measurements were performed in a blind manner, and then groups of different Gleason grades were made for significance analysis.

For fractal dimension calculation, each image is used as a color channel split for distinguishing cells, stroma and lumen. In this case it was found that the blue color filter differentiated the stroma, the green channel the lumen, while the red color filter differentiated the cell nuclei, as shown in Figure 2. Each filtered image was converted into binary and the fractal dimension was determined by the box count method [23].

**Figure 2.**
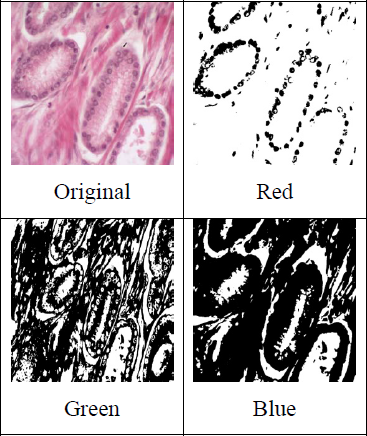
Filtering color process of a image of a prostate cancer histopathological section.

The *LZ* complexity [24, 25] was implemented as follows: the histopathological images are represented by a matrix [I (X, Y)], where each coordinate (X, Y) represents a pixel of the image, or more properly a intensity value of one color (Fig. 2). This way discrete series in coordinates “X” and “Y” respectively is generated to calculate the *LZ* complexity thereof. It is considered that each row of the image (X1,2,…, width) is a time series and its *LZ* complexity is calculated, (LZ_X) and the same way for each column (Y1,2,…, height) (LZ_Y). Accordingly, if the image is (1296×972 pixels) we have 1296 rows and 972 columns and thus following this idea, we have two time series: one consisting of 1296 LZ_X values and other of 972 LZ_Y values. In this way the image is reduced to a time series of complexity in X and Y (Fig. 3). To these new time series the *LZ* complexity is calculated once again to reduce to a single value in X (LZ_X_last) and Y (LZ_Y_last).

To asses significant differences of the different Gleason results for fractal dimension and *LZ* complexity, a t-test for two independent populations at a level of significance of *p*<0.05.

**Figure 3.**
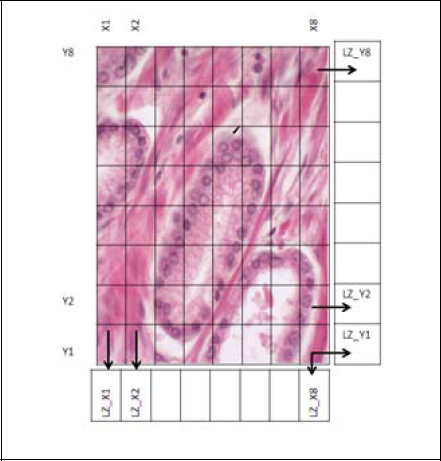
Procedure to get values of *LZ* complexity.

## 3. Results

Figure 4 shows histopathological images for increasing Gleason grades, indicative of an increase in the tumor agresiveness. Gland morphology progressively modified, observed as loss of the auto-organized structure of glands and decrease of the area occupied by the lumen (Figure 4).

**Figure 4.**
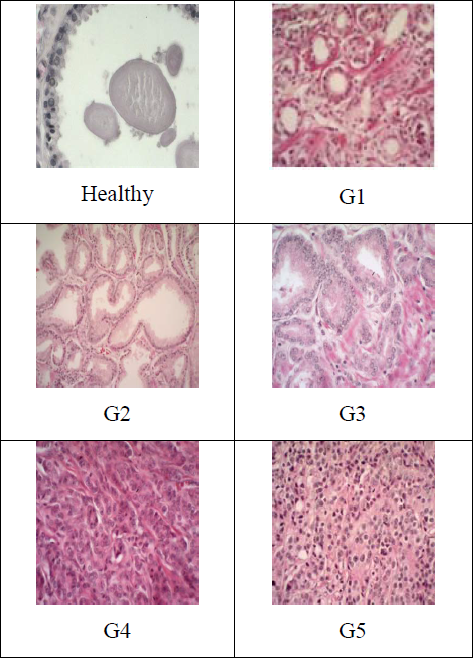
Distribution of tissue pattern in prostate cancer to different Gleason grades.

We determined the fractal dimension on the lumen of each image and the *LZ* complexity of all histopathological images (Table 1). Comparison between groups using the t-test (Table 2) shows that *p* values for 60% of cases is less than 0.05 (*p*<0.05) indicating that there are significant differences in the fractal dimension of the lumen and the *LZ* complexity of the prostatic tissue between healthy tissue and tumor at the different Gleason grades.

**Table 1.** Average values of *LZ* Complexity and *D_f_* of lumen of healthy tissue and tumor prostate of increasing Gleason grade.

| Histology | $D_f$ (Lumen) | $LZ$ | N |
| --- | --- | --- | --- |
| Normal | 1.71±0.02 | 0.52±0.03 | 30 |
| Gleason 1 | 1.54±0.03 | 0.77±0.03 | 5 |
| Gleason 2 | 1.64±0.03 | 0.68±0.03 | 12 |
| Gleason 3 | 1.58±0.01 | 0.65±0.01 | 76 |
| Gleason 4 | 1.46±0.03 | 0.67±0.03 | 15 |
| Gleason 5 | 1.48±0.03 | 0.79±0.02 | 6 |

**Differences between grades specimens are statistically significant ( $p<0.05$ )**

**Table 2.** t-Test at significance level *p* =0.05 obtained for *LZ* Complexity and *D_f_* of lumen of healthy tissue and tumor prostate of increasing Gleason grade.

| $LZ$ | | | | | |
| --- | --- | --- | --- | --- | --- |
|  | Healthy | G1 | G2 | G3 | G4 |
| G1 | 0.00 |  |  |  |  |
| G2 | 0.00 | 0.09 |  |  |  |
| G3 | 0.00 | 0.02 | 0.34 |  |  |
| G4 | 0.00 | 0.09 | 0.73 | 0.59 |  |
| G5 | 0.00 | 0.45 | 0.02 | 0.00 | 0.02 |

| $D_f$ | | | | | |
| --- | --- | --- | --- | --- | --- |
|  | Healthy | G1 | G2 | G3 | G4 |
| G1 | 0.00 |  |  |  |  |
| G2 | 0.05 | 0.06 |  |  |  |
| G3 | 0.00 | 0.34 | 0.09 |  |  |
| G4 | 0.00 | 0.15 | 0.00 | 0.00 |  |
| G5 | 0.00 | 0.21 | 0.00 | 0.01 | 0.70 |

As expected, the fractal dimension (*D_f_*) of lumen of healthy tissue and tumor prostate of increasing Gleason grade was progressively decreasing, exhibiting an acceptable correlation (Figure 5).

**Figure 5.**
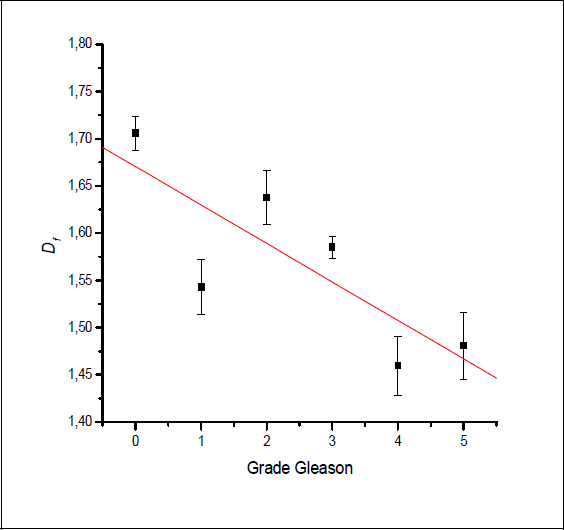
Average and SE values of *D_f_* for each Gleason grade. Healthy tissue is considered as grade 0.

Interestingly, *LZ* complexity allows to istinguish between healthy and prostate cancer images (Figure 6). In addition, *LZ* complexity first decreases and later increases with increasing agressiveness of prostate cancer. This shift in the curve tendency occurs at Gleason grade 3 (Figure 6).

**Figure 6.**
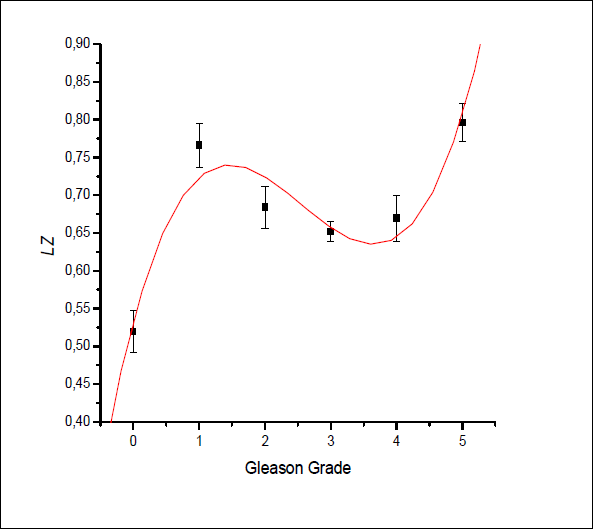
Average and SE values of *LZ* for each Gleason grade. Healthy tissue is considered as grade 0.

## 4. Discussion

Recent studies [20, 21], set the fractal dimension as an additional histopathological analysis method for prostate cancer, but these authors only set guidelines to separate malignant and benign prostate tissue, without deepen in the real problem of the standardization of malignancy grades classified according to the Gleason grade.

As shown in Figure 5, there is an acceptable correlation between the fractal dimension of the lumen and the tumor degree. The experimental results obtained show the loss of a gland self-organized structure of the histopathological patterns from Gleason 1 onwards, suggesting an increase in the tumor aggressiveness with the decrease of the fractal dimension. This is a mathematical validation of the value of the score that Gleason established more than 50 years ago [9].

The dynamic of *LZ* shown in Figure 6 exhibits an abrupt change in slope in the vicinity of grade 3. Mathematically it behaves like a phase transition of the first order [26]. This is an important indicator of the changes observed by pathologists in this grade in this particular type of cancer. In fact, up until Gleason grade 3, the glandular-like structure is conserved, and at grades 4 and 5 is lost. This mathematical data provides validation for the consensus of the ISUP considering the poorly formed glands and cribiform pattern to be higher than the original Gleason 3 (poorly formed glands and cribiform pattern are now considered Gleason 4) [10, 11].

Moreover, the determination of the *LZ* complexity of the histological pattern, has the advantage that, unlike the fractal dimension, the determination is done considering the histopathological image as a whole.

The results show that it is possible to characterize the complexity of the tissues of prostate cancer through the values of fractal dimension of the lumen *D_f_* and *LZ* complexity. Thereby the results obtained are a quantitative complement to diagnosis and consequently could contribute to the development of new strategies and improved prognosis with future potential automation of the method. This method allows the pathologist to suggest the therapeutic strategy more adequate for each patient considering the tumor aggressiveness, allowing for a more accurate personalized medicine treatment.

In conclusion, we propose to quantify the prostate tumor agressiveness through the fractal dimension of the lumen and the *LZ* complexity of the image as a whole. The fractal dimension and *LZ* complexity provide a complementary quantitative characterization of the different grades of the tumor, which is a nominal variable, and able to differentiate healthy tissue.

## Acknowledgements

This work was support in part by Spanish Agency of International Cooperationfor Development (AECID, projects: D/023653/09 and D/030752/10), the project SAF2010-19230 of Ministerio Español de Ciencia e Innovación to the Cell engineering Laboratory, IdiPAZ, as well as Biomedicine and Biotechnology, University of Alcalá; and Mexican Institute of Complex Systems, Geo-estratos S.A., México.

